# Statistical mechanics of cell decision-making: the cell migration force distribution

**DOI:** 10.1101/235689

**Authors:** Haralampos Hatzikirou

## Abstract

Cell decision-making is the cellular process of responding to microenvironmental cues. This can be regarded as the regulation of cell’s intrinsic variables to extrinsic stimuli. Currently, little is known about the principles dictating cell decision-making. Regarding cells as Bayesian decision-makers under energetic constraints, I postulate the principle of least microenvironmental uncertainty (LEUP). This is translated into a free-energy principle and I develop a statistical mechanics theory for cell decision-making. I exhibit the potential of LEUP in the case of cell migration. In particular, I calculate the dependence of cell locomotion force on the steady state distribution of adhesion receptors. Finally, the associated migration velocity allows for the reproduction of the cell anomalous diffusion, as observed in cell culture experiments.

## I. INTRODUCTION

Cell decisions are responses of the cell’s intrinsic mechanisms to extrinsic signals [1]. Such mechanisms are typically termed as signal transduction pathways. Typically, signal transduction occurs when an extracellular signalling molecule binds to a certain receptor creating a complex. In turn, this complex of molecules triggers a biochemical chain of events inside the cell [2]. These biochemical events can influence the epigenetic or even the genetic state of the cell. Then the new genomic state of the cell will be reflected in cell’s transcriptome and translated into proteins that generate a phenotypic response. In general, cells encode extrinsic information into new intrinsic states and in turn decode the latter into phenotypic responses.

Concerning cell decision-making there are two fundamental issues: (i) typically there is a great deal of uncertainty in the intracellular dynamics involved in cell phenotypic decisions, and (ii) although there is a lot of work focused on single cell cell decision-making [] in the context of multicellular systems little is known. In particular, there are two schools of thought to address the single cell decision-making.

The first school thought is a *mechanism-driven* approach which follows the idea of epigenetic landscape reconstruction [3]. This dates back to Waddigton epigenetic landscape where cell phenotypes trickle down along a bifurcating landscape of “valleys” and “hills” (see Fig. 1). Reconstructing such landscapes requires very good knowledge of the underlying genetic, epigenetic or transcriptional dynamics and provides elegant mathematical formulations that allow for temporal predictions. Obviously, this approach suffers from the issue (i), i.e. we do not know all possible intracellular players and dynamics involved in phenotypic decision.

**Fig. 1.**
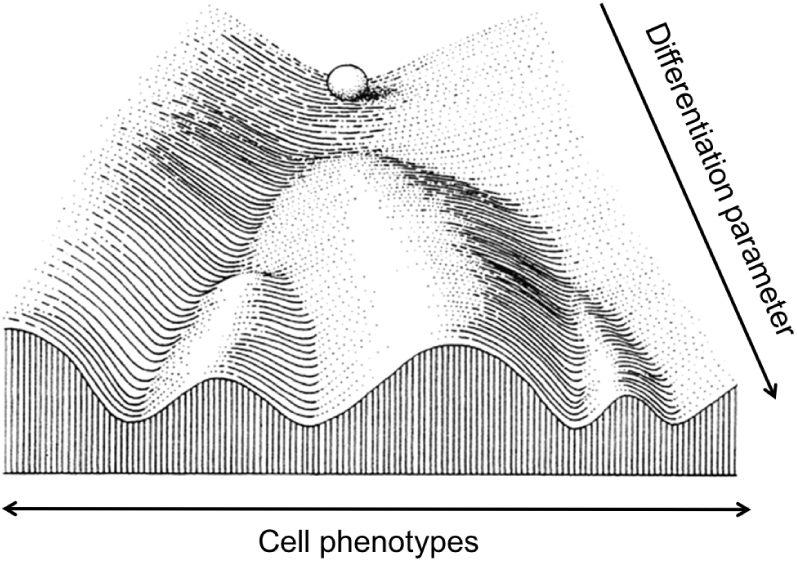
Sketch of communication between cell and its microenvironment

The second is a *data-driven* approach involving ideas from statistical inference [4]. In principle, cell decisions are treated in a mechanism-blind way, as an input-output system, without requiring the knowledge of the intracellular components involved in phenotypic regulation.The intracellular dynamics are treated as a black box. The main goal is to optimize quantities such as mutual information and infer the corresponding phenotypic responses. The main problem with this approach is that ignores dynamics, i.e. does not allow for temporal predictions or analysis of the cells phenotypic dynamics.

Given the above problems, my goal is to identify a variational principle for cell decision-making that fulfils the following: (i) keeps the elegance and predictability of a dynamic description, and (ii) circumvent the details of intracellular dynamics as statistical inference. Here I propose the least microenvironmental uncertainty principle (LEUP) which I develop further in a statistical mechanics theory for cell decision making. Finally, I apply the LEUP theory into the cell migration problem and I calculate the corresponding force distribution and migration velocity.

## II. THE BAYESIAN CELL DECISION-MAKER

Cells typically collect information from their microenvironment and encode it into tangible actions, i.e. phenotypic decisions. The latter involves the rearrangement of their intrinsic variables x ∈ X ⊂ ℝ^n^, representing genes, RNA molecules, translational proteins, metabolites, membrane receptors, etc. Microenvironmental or extrinsic variables are denoted by y ∈ Y ⊂ ℝ^n^ being ligand, chemicals, nutrients, cellular density or stress fields. In order to formalise this encoding process, we consider a Bayesian cell decision making:

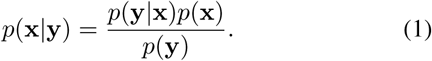

In particular, the extrinsic variable distribution *p*(y|x) represents the likelihood of mircoenvironmental information being collected by the cell. In turn, it is combined with the prior probability distribution function (pdf) of intrinsic variables *p*(x) resulting into the posterior intrinsic variable distribution *p*(x|y). The latter represents the decision of the cell over the available information.

The collection of microenvironmental data, i.e. assesing the likelihood *p*(y|x), is energetically expensive. For instance, as shown in Fig. 2, sampling the cellular microenvironment involves the recruitment/degradation of different receptors (intrinsic state) and the corresponding binding to diffusible ligands (extrinsic state). As any physical system, cells would try to minimize energetic costs unless otherwise required. Therefore, I postulate that an energy-efficient cell would require the construction of the least informed/optimal prior^1^. This idea is rather straight forward in the case of differentiated cells, since heart cells, for instance, sufficiently know their heart microenvironment and they would fail living in an other one, e.g. skin.

**Fig. 2.**
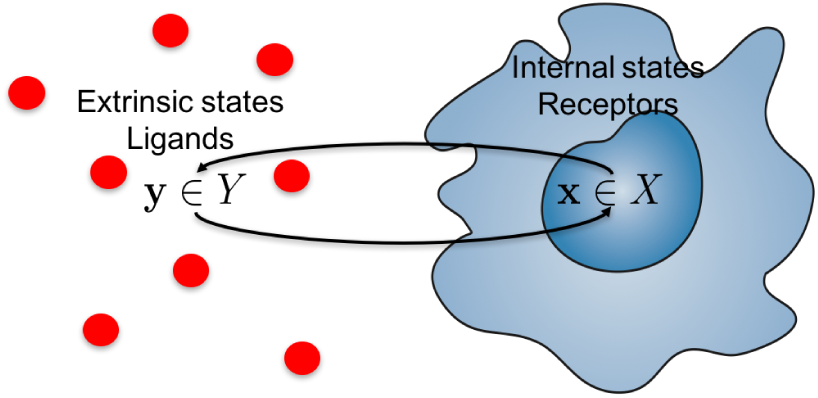
Sketch of communication between cell and its microenvironment

## III. A FREE-ENERGY PRINCIPLE: THE LEAST MICROENVIRONMENTAL UNCERTAINTY PRIOR

The above arguments could be translated in a statistical mechanics formulation. My central idea is that the cell decides on its phenotype by virtue of minimally sampling its microenvironment. As stated above, my goal is to optimize the cell’s prior and at the same time minimize the total uncertainty of the system cell-microenvironment. This can be mathematically translated into finding the appropriate intrinsic state pdf *p*(x) that minimizes the joint entropy of cell intrinsic and extrinsic variables

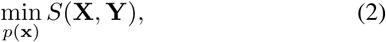

where 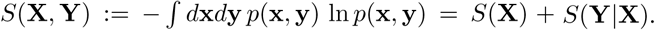

The corresponding variational principle that follows reads:

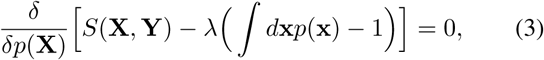

where the 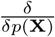 denotes the functional derivative operator and λ a Lagrange multiplier for the normalization constraint. The solution of eq. (3) is the equilibrium pdf:

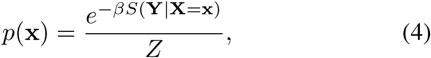

where 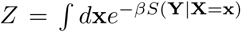 the corresponding normalization factor. The last equation is very interesting since it shows the equilibrium distribution of the cell’s intrinsic states is the one that confers the least uncertainty of microenvironmental conditions. Please note that I introduced the parameter *β* as an “inverse temperature”, that quantifies the compliance of a cell to LEUP. Please note that additional biological knowledge will be translated into Lagrange constraints of the free-energy minimisation.

To illustrate the above result, let us assume that the distribution *p*(y|x) follows the normal distribution 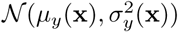. Then the mutual entropy reads

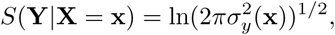

and the eq. (4) respectively becomes:

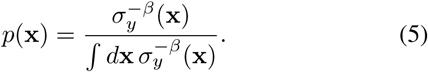

This last equation is exactly the normalized square root of Fisher’s Information for the second moment, i.e. 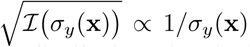 [5]. Given the latter our equilibrium distribution (5) coincides with a normalized Jeffrey’s prior distribution, for *β* = 1. Jeffrey’s prior represents the definition of non-informative Bayesian prior in classical statistics [6].

Substituting the equilibrium pdf (4) into the cell intrinsic state entropy, we obtain:

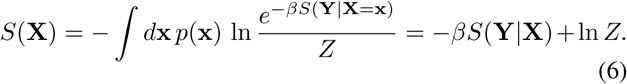

The above is a fundamental thermodynamic identity of LEUP where the first term denotes the system’s total phenotypic internal energy 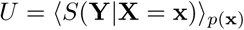 which is the average microenvironmental uncertainty sensed by a cell. The last term represent a phenotypic free energy *F* = *β*^−1^ ln *Z*.

## IV. CELL MIGRATION FORCE PROBABILITY DISTRIBUTION AGAINST INTEGRIN RECEPTORS

To illustrate the implications of the LEUP in a single cell-microenvironment system, we assume the problem of single cell migration (see Fig. 3). Cell membrane-expressed integrin receptors bind to the corresponding extra-cellular matrix (ECM) ligands. The formation of stable focal adhesion points (FA), i.e. bond clusters, is required to generate locomotion force. Cell migration starts when the force at the front of the cell is larger than the force at the rear, inducing the rupture of the rear cell bonds. Integrin receptors play the role of cell’s intrinsic state and the a density of ECM ligands the corresponding microenvironmental/extrinsic variable [7]. Here, I try to calculate the locomotion force distribution via the surface-expressing receptor distribution.

**Fig. 3.**
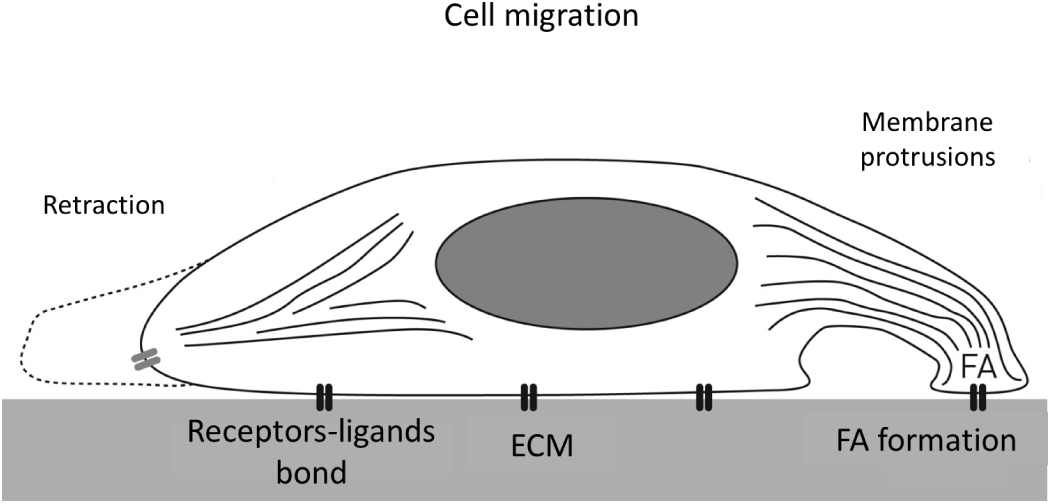
Simplified description of cell migration processes.

I employ the simplest FA dynamics model of receptor-ligand concentrations denoted by *x*(*t*) and *y*(*t*), respectively, and *c*(*t*) corresponding to bond density:

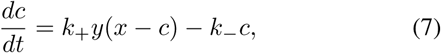

where *k*_+_ the association and *k*_-_ the disassociation rate. Then, I calculate the steady state value of bond concentration:

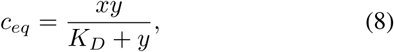

where the parameter *K*_*D*_ = *k*_-_/*k*_+_. It is well-known that the disassociation depends on the adhesion force *F* as *k*_-_ = *e*^*F/c*^ and in the steady state one can prove (for details see in Schwartz et al [7]) that:

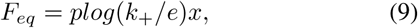

where the function *plog* is the solution of the equation *xe^x^* = *a*. Therefore, the equilibrium force pdf is proportional to the steady state distribution of the receptors.

At this point, in order to calculate the equilibrium probability of receptors *p*(*x*), I need to estimate the bond fluctuations of the system. The fluctuations estimated at equilibrium read:

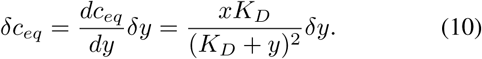

Using the Fourier transform 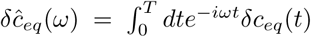, we can state that the standard deviation of *c_eq_* is related to the power spectral density 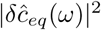 as

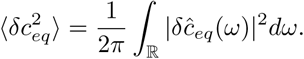

The next goal is to calculate the steady state variation of complex formation. In this respect, we use the master equation as described in [8]:

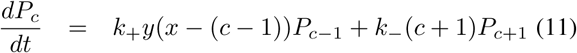

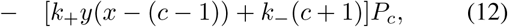

where *P_c_*(*t*) the probability distribution of complexes and *c* = 1,2, …, *x* — 1. The resulting probability generating function reads at the steady state:

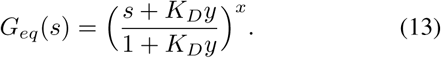

Then the complex distribution coincides with the Binomial one with parameter (1 + *K_Dy_*)^−1^. The variance is simply given by the following formula:

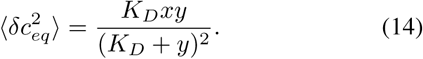

Combining equations (10) and (14), we can obtain the steady state fluctuations of the ligand density:

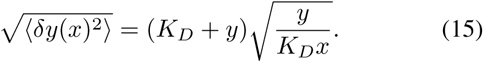

Finally, assuming that ligands for a given receptor concentration follow a normal distribution and a “inverse temperature” *β*, the equilibrium pdf of receptor is provided by eq. (5):

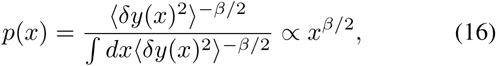

which results into power law distributions for different values of *β*. Assuming normality of the ligand *y* distribution and *c_eq_* (the binomial distribution converges to the normal one for large numbers of receptors), then from the eq. (8) and for *K_D_* ≫ 1, we have *x* α *c_eq_*/*y*. One can easily prove that the ratio of two Gaussian distributions results into fat-tailed Cauchy distribution which scales the data, i.e. 1/*x^2^*. LEUP would fit the corresponding data for *β* = —4.

The above result states that the cell locomotion force pdf is proportional to the power law distribution of the adhesion receptors

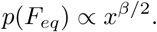

In particular it implies that there is a finite probability for cells to exert high locomotion forces. In principle, the presence of power laws would support the long-standing hypothesis that biology is poised at criticality [9].

It is very interesting to see the impact on the cell locomotion velocity, where using Newton’s law the cell velocity reads:

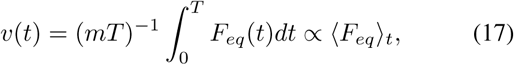

where *m* the cell mass and *T* the total time. Assuming ergodicity in the force generation process, then the above temporal average is equal to the ensemble average, i.e. ⟨*F_eq_*⟩_*t*_ = ⟨*F_eq_*⟩ _*F*_. Thus the cell velocity reads, using eqs (9,16):

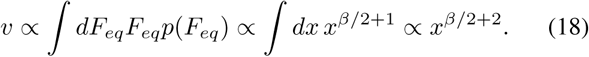

In the same result we could arrive by assuming that cells operate in the overdamped limit.

Now assuming that *x* ∝ *t^a^*, where *a* ≥ 0, one recovers the superdiffusive cell migration dynamics as observed in cell motility experiments [10] for *β >* —4 (see Fig. 4), since

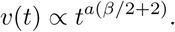

**Fig. 4.**
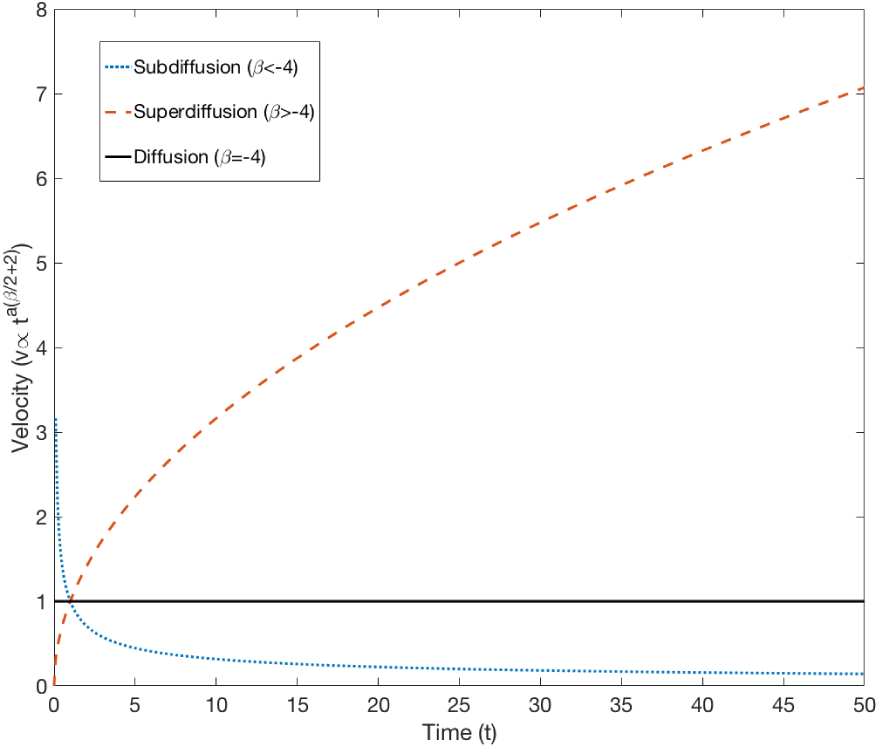
The LEUP compliance *β* controls the different regimes of cell diffusion.

In the low association limit *K_D_* ≫ 1, the LEUP compliance was estimated as *β* = —4. Then the average cell velocity is equal to a constant, i.e. the cell diffuses randomly. This is rather expected, since no stable focal adhesion points can be established and the cell cannot establish a persistent migration direction. From the inverse point of view, random motion is not expected to decrease local microenvironmental uncertainty as LEUP dictates, therefore we could expect a negative *β*.

## V. DISCUSSION

In this article, I present a novel statistical theory for cell decision-making, the so-called least microenvironmental uncertainty principle (LEUP). This is translated into a variational principle that minimized the joint entropy of cell intrinsic states and the corresponding microenvironental states. In turn, I develop the statistical mechanics theory for equilibrium cell decision-making, where I derive thermodynamic-like relationships. In turn, I apply the theory in a simplified model of cell migration and I calculate the locomotion force distribution. Finally, the resulting force distribution recovers the observed single cell anomalous diffusion.

In the present paper, I develop only the equilibrium theory. However, the minimization of the LEUP free energy allows us to formulate cell decision-making dynamics. This can be in the form of Monte-Carlo simulations or Langevin-like stochastic differential equations. In a future work, I will elaborate on this direction.

One could note that this is not the first effort to use information theoretic arguments to understand cell fate determination. Similar approaches have been undertaken by other works, such as [11], [12], [13], [14]. However, in another paper I discuss the connection of all these approaches with LEUP [15].

Finally, LEUP is particular interest since it allows for the inference of cellular intrinsic states/phenotypes by means of the local microenvironental entropy or fluctuations. This allows for computation of the cellular states without the explicit understanding the underlying mechanisms. The sole knowledge of the extrinsic variable distribution is enough. Therefore, we could apply the LEUP to problems where phenotypic regulation is unclear or unknown. One direct application would be on the regulation of migration/proliferation plasticity or Go-or-Grow in the context of glioblastoma tumors [16], [17].

## Footnotes

1 An informed prior is the one that requires the most sampling of new data. Here we are interested to least collection of data, i.e. the least informed prior.

